# TFEB/HLH-30-mediated expansion of neuronal lysosomal capacity in early adulthood protects dendrite maintenance during aging in *Caenorhabditis elegans*

**DOI:** 10.1101/2024.11.29.625995

**Authors:** Ruiling Zhong, Claire E. Richardson

## Abstract

Lysosomes are essential for neuronal homeostasis, providing degradation and recycling functions necessary to support neurons’ complex operations and long lifespans. However, the regulation of lysosomal degradative capacity in healthy neurons is poorly understood. Here, we investigate the role of HLH-30, the sole *Caenorhabditis elegans* homolog of Transcription Factor EB (TFEB), a master regulator of lysosome biogenesis and autophagy that it is thought to predominantly function in the context of starvation or stress. We demonstrate that HLH-30 is dispensable for neuronal development but acts cell-intrinsically to expand lysosomal degradative capacity during early adulthood. Loss of HLH-30 leads to lysosomal dysfunction and delayed turnover of synaptic vesicle proteins from the synapse. Notably, we show that basal HLH-30 activity is sufficient to expand neuronal lysosomal capacity without nuclear enrichment, in contrast to the nuclear translocation associated with starvation- and stress-induced activation of TFEB and HLH-30. Furthermore, we show that neuronal lysosomal function declines with age in wild-type animals, and this corresponds to a decrease in basal HLH-30-mediated transcription. We further demonstrate that basal HLH-30 activity is crucial for neuron maintenance: lysosomal dysfunction due to inadequate HLH-30 activity leads to dendrite degeneration and aberrant outgrowths. In summary, our study establishes a critical role for HLH-30/TFEB in promoting lysosomal capacity to preserve neuronal homeostasis and structural integrity of mature neurons *in vivo*.

## Introduction

To sustain their long lifespans, complex morphologies, and diverse functions, neurons require robust mechanisms of cellular homeostasis. Lysosomes are key effectors of cellular homeostasis. Using the dozens of enzymes within their acidic lumens, lysosomes degrade transmembrane proteins, protein aggregates, and organelles, promoting their proper abundance and quality control. In neurons, degradative lysosomes reside predominantly in the cell body [1–3]. Their cargoes are earmarked for degradation throughout the neuron primarily via the endolysosomal pathway, which sorts transmembrane proteins for degradation into late endosomes, or through autophagy, in which autophagosomes engulf organelles and protein aggregates [2,4–8]. Once transported to the cell body, these compartments fuse with lysosomes to generate endolysosomes and autolysosomes, respectively, where the cargo is degraded before the lysosomes reform, ready for the next load [4,9,10]. For neurons to maintain effective lysosomal degradation, their lysosomal capacity – the number and functionality of lysosomes – must be sufficient to meet their high demands for cellular turnover. Yet, despite its importance, how neuronal lysosomal capacity is regulated remains largely unknown.

A potential regulator for neuronal lysosomal capacity is Transcription Factor EB (TFEB), which regulates the expression of genes involved in autophagy and lysosome biogenesis [11–14]. The activity of TFEB is best understood in the context of nutrient sensing: in nutrient-rich conditions, the mammalian target of rapamycin complex 1 (mTORC1) phosphorylates TFEB, leading to its cytosolic retention and inactivation. During starvation, mTORC1 activity is inhibited, allowing dephosphorylated TFEB enter the nucleus to induce the expression of endosomal, lysosomal, and autophagosomal genes [15–17]. Similarly, HLH-30, the sole *C. elegans* homolog of TFEB, regulates lysosome biogenesis and autophagy genes in response to stress [18,19]. Like mammalian TFEB, HLH-30 predominantly localizes to the cytoplasm under well-fed conditions [11,13,19]. However, starvation or various stressors promotes its nuclear translocation in hypodermal and intestinal cells [20–22]. The role of TFEB/HLH-30 in neurons under non-stressed conditions remains largely unexplored.

Understanding the regulation of neuronal lysosomal capacity has important health implications, as declining endolysosomal and autophagic function is closely linked to neurodegenerative diseases [23–28]. Lysosomal dysfunction plays a central role in neurodegenerative diseases, where protein aggregates, which are normally targeted for lysosomal degradation, accumulate and cause cellular toxicity [29–31]. There is evidence that lysosomal dysfunction is also a pathology of normal aging, leading to speculation that age-associated lysosomal dysfunction in neurons may contribute to neurodegenerative disease. For example, lipofuscin - an autofluorescent mix of oxidized macromolecules – accumulates within neuronal lysosomes during aging, which is thought to be indicative of and/or cause impaired lysosomal function [32,33]. Aging has also been associated with the accumulation of late endosomes in the presynapses in both mouse and *Drosophila* neuromuscular junctions, which could arise from an impairment in lysosomal function [34,35]. A reduced fusion rate between autophagosomes and lysosomes, along with the failure to deliver autophagosome substrates into lysosomes in aged mice, indicates that neuronal autophagy declines with aging [36,37]. Furthermore, axonal transport of autophagosomes and late endosomes, which relies on acidification from fusing with lysosomes, show defects in aging mice [38–42]. This evidence highlights the need for a better understanding of whether and how neuronal lysosomal capacity becomes inadequate in healthy aging, to inform interventions against neurodegenerative disease.

Here, we first present direct evidence that neuronal lysosomal function declines during healthy aging *in vivo*. Next, we showed that HLH-30 functions neuron-intrinsically to promote adequate lysosomal capacity in adulthood. Loss of HLH-30 accelerates the age-related decline in lysosomal function, and it causes delayed degradation of typical cargoes for endolysosomal degradation – synaptic vesicle proteins – leading to a backlog of degradative materials at the synapse. Interestingly, we found that basal HLH-30 activity supports neuronal lysosomal capacity without enriched nuclear localization, in contrasts to the nuclear enrichment associated with TFEB and HLH-30 activity during starvation and stress. Additionally, HLH-30-regulated gene expression shows a declining trend with age, suggesting that reduced HLH-30 activity contributes to age-associated lysosomal dysfunction. Finally, we demonstrated that neuronal HLH-30 is critical for neuronal maintenance, as its loss accelerates dendrite age-related degeneration and aberrant dendritic sprouting. Based on these results, we propose that TFEB/HLH-30 performs a crucial role in promoting lysosomal capacity to preserve neuronal homeostasis during adulthood *in vivo*.

## Results

### Lysosome function declines during aging in *C. elegans* neurons

To test the hypothesis that neuronal lysosomal dysfunction is a pathology of aging *in vivo*, we examined the posterior ventral process D (PVD) neurons. These two bilaterally symmetric, glutamatergic sensory neurons provide an accessible model to interrogate lysosomal function with high resolution. Lysosomal degradation occurs predominantly in the neuron cell body, and as in other organisms, the lysosomal compartments within *C. elegans* neurons mostly localize to the cell body [43]. We therefore focused on the PVD neuron cell body and assessed three established features of lysosomal dysfunction: increased luminal pH, lysosomal compartment enlargement, and buildup of degradative cargo within lysosomal compartments [44].

First, we developed a tool to measure relative lysosome acidity. Healthy lysosomes have a luminal pH of about 5, and a higher pH indicates lysosomal dysfunction [3,45]. NUC-1, the ortholog of human DNAse II, is a luminal lysosome resident and a previously validated marker for lysosomal compartments – lysosomes, endolysosomes, and autolysosomes – in *C. elegans* [46]. We co-expressed NUC-1::GFP and NUC-1::RFP alongside a BFP morphology marker in PVD neurons (Fig 1A, S1A Fig). GFP fluorescence is partially quenched in the acidic environment of lysosomal compartments, while RFP intensity is unaffected by the lysosomal pH and is used for normalization. Therefore, the GFP/RFP ratio represents relative acidity: a higher ratio indicates reduced acidity (i.e., higher pH). We first validated our tool using wild-type (WT) worms treated with 0.03 μM Concanamycin A, which inhibits vacuolar H^+^ ATPase. Relative lysosomal acidity is decreased in Day 6 adults post-treatment compared to untreated animals, demonstrating the serviceability of our tool (Fig 1B). We then examined WT worms at different ages and found that relative neuron lysosome acidity decreases in aged worms by Day 9 of adulthood. For comparison, the median lifespan of worms is about two weeks [19,47–49].

**Fig 1.**
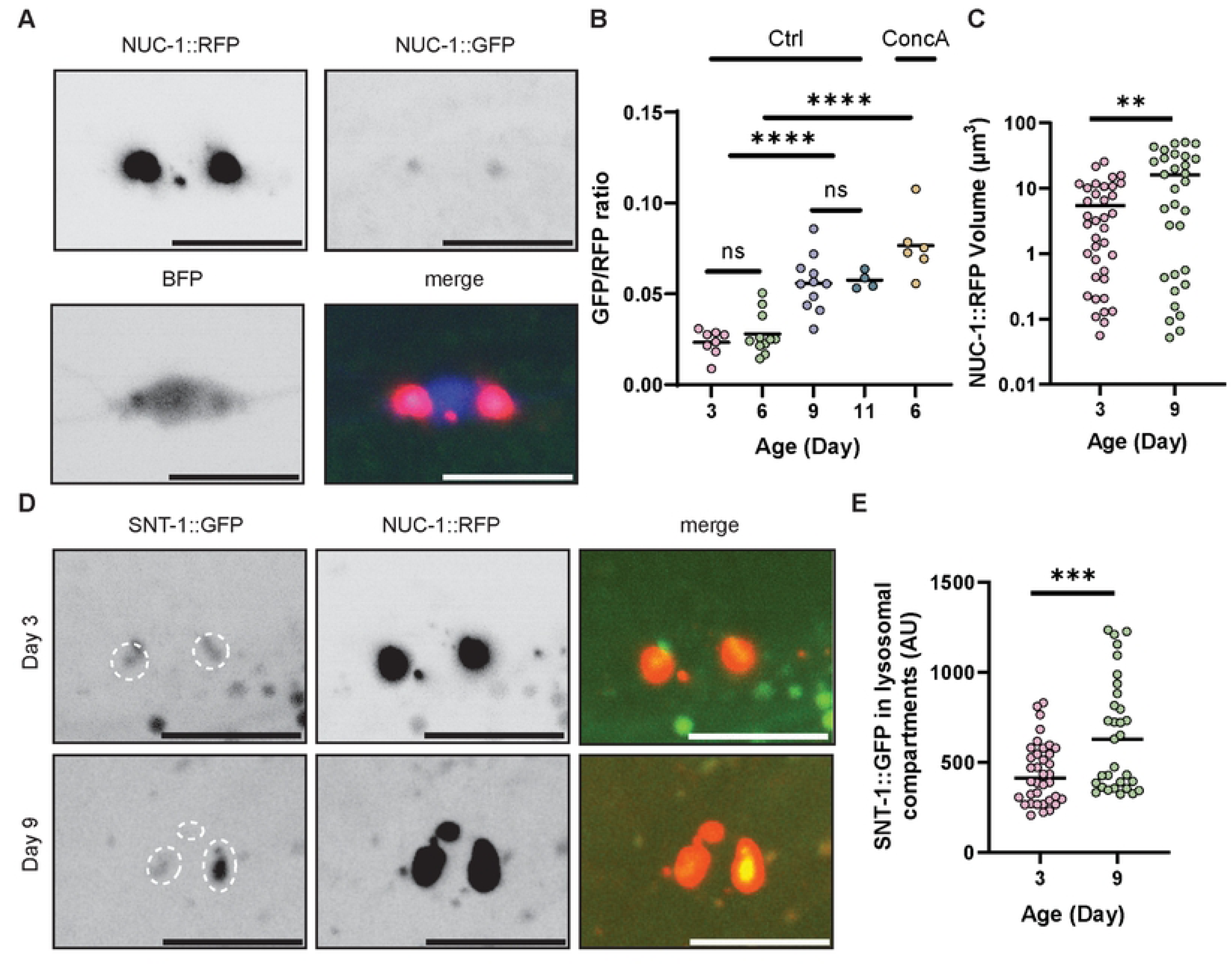
PVD neurons show reduced lysosome function during aging. (A-B) A fluorescent reporter for luminal acidity of neuronal lysosomal compartments shows that acidity decreases with aging. (A) Example images of the PVD neuron cell body in a Day 3 adult carrying the *carEx6* transgene, which stably expresses all three constructs shown. (B) Quantification of NUC-1::GFP/RFP ratio using the *carEx6* transgene. Scale bar = 10 μm, **** P < 0.0001, ns: not significant, One-way ANOVA with Tukey post-test. ConcA: Concanamycin A. (C) Aged animals have larger lysosomal compartments than younger adults. **P < 0.01, Kolmogorov-Smirnov test. (D-E) A representative degradative cargo for lysosomes, the synaptic vesicle transmembrane protein SNT-1, shows increased accumulation within lysosomal compartments in older adults compared to younger adults. Scale bar = 10 μm. ***P < 0.001, two-tailed *t* test.

Second, we quantified lysosomal compartment size using NUC-1::RFP. At Day 3 of adulthood, we observed a wide range of lysosomal compartment sizes (Fig 1C). This reflects lysosomal compartment dynamics: in the neuron cell body at any moment, we expect to observe tiny lysosomes as well as the larger endolysosomes and autolysosomes. By Day 9 of adulthood, the mean size of NUC-1::RFP cell body puncta increased compared to Day 3 (Fig 1C). This difference is driven by the larger size of the largest lysosomal compartments in Day 9 adults. The smallest puncta, which likely represent true lysosomes, show a similar size between Day 3 and Day 9 animals. We note that with the “large” lysosomal compartments, we cannot resolve which puncta are individual lysosomal compartments versus clusters of multiple smaller lysosomal compartments using light microscopy; still, both enlargement and clustering indicate lysosome dysfunction [46,50,51].

Third, we assessed the efficiency of lysosomal degradation by quantifying the accumulation of a degradative cargo, the synaptic vesicle protein Synaptotagmin/SNT-1, within lysosomal compartments in the neuron cell body. We measured the fluorescence intensity of endogenously-tagged SNT-1::GFP that co-localized with NUC-1::RFP, which represents SNT-1 protein that has been sorted into a lysosomal compartment but has not been fully degraded, and the GFP has not been fully quenched (Fig 1D, S1B Fig). By Day 9 of adulthood, the intensity of SNT-1::GFP in lysosomal compartments was higher than at Day 3, supporting the notion of age-related lysosomal dysfunction (Fig 1E). Taken together, these data support the model that neuronal lysosomal dysfunction is a pathology of aging *in vivo*.

### HLH-30 is required for adult neuronal lysosomal capacity and function

To investigate the cause of declining neuronal lysosomal function in aged animals, we first need to understand how neurons maintain adequate lysosomal function in early adulthood.

TFEB is a master regulator of lysosome and autophagosome biogenesis in mammalian cells, but it has been predominantly shown to function during starvation and stress [52]. Whether and how TFEB functions in healthy neurons *in vivo* is unknown. We first asked whether loss of *hlh-30*, the *C. elegans* homolog of *TFEB*, affects the late endosome, endolysosome, and lysosome population in the neuron. The *hlh-30(o)* mutant is viable and displays WT animal morphology and behavior [19]. Likewise, the morphology of the PVD neuron in the *hlh-30(o)* mutant appears indistinguishable from WT (S2 Fig). The RAB-7 GTPase is a trafficking factor of late endosomes and autophagosomes, and it localizes to late endosomes, endolysosomes, and autolysosomes [38]. Using PVD-expressed mCherry::RAB-7, we found that the number of RAB-7-labeled compartments in the neuron cell body increases between Day 0 of adulthood (the L4 larval stage) to Day 3 of adulthood (Figs 2A-B) [53]. In the *hlh-30(o)* mutants, the number of RAB-7 labeled compartments is no different at Day 0 but significantly reduced at Day 3 of adulthood compared to WT (Fig 2B). This reduction at Day 3 can be rescued by expressing *hlh-30::gfp* specifically in the PVD neurons, indicating that HLH-30 regulates the lysosomal compartment abundance cell-autonomously. We observed the same trends when we quantified the abundance of endogenously-tagged GFP::RAB-7 compartments in WT versus *hlh-30(o)* mutants, indicating that the accumulation of RAB-7-labeled lysosomal compartments in adult worms and their reduced abundance in *hlh-30(o)* mutants are not due to an overexpression artifact (S3A Fig) [53]. These data suggest that HLH-30 promotes late endosome, endolysosome, and/or autophagosome abundance in early adulthood but is dispensable in development.

**Fig 2.**
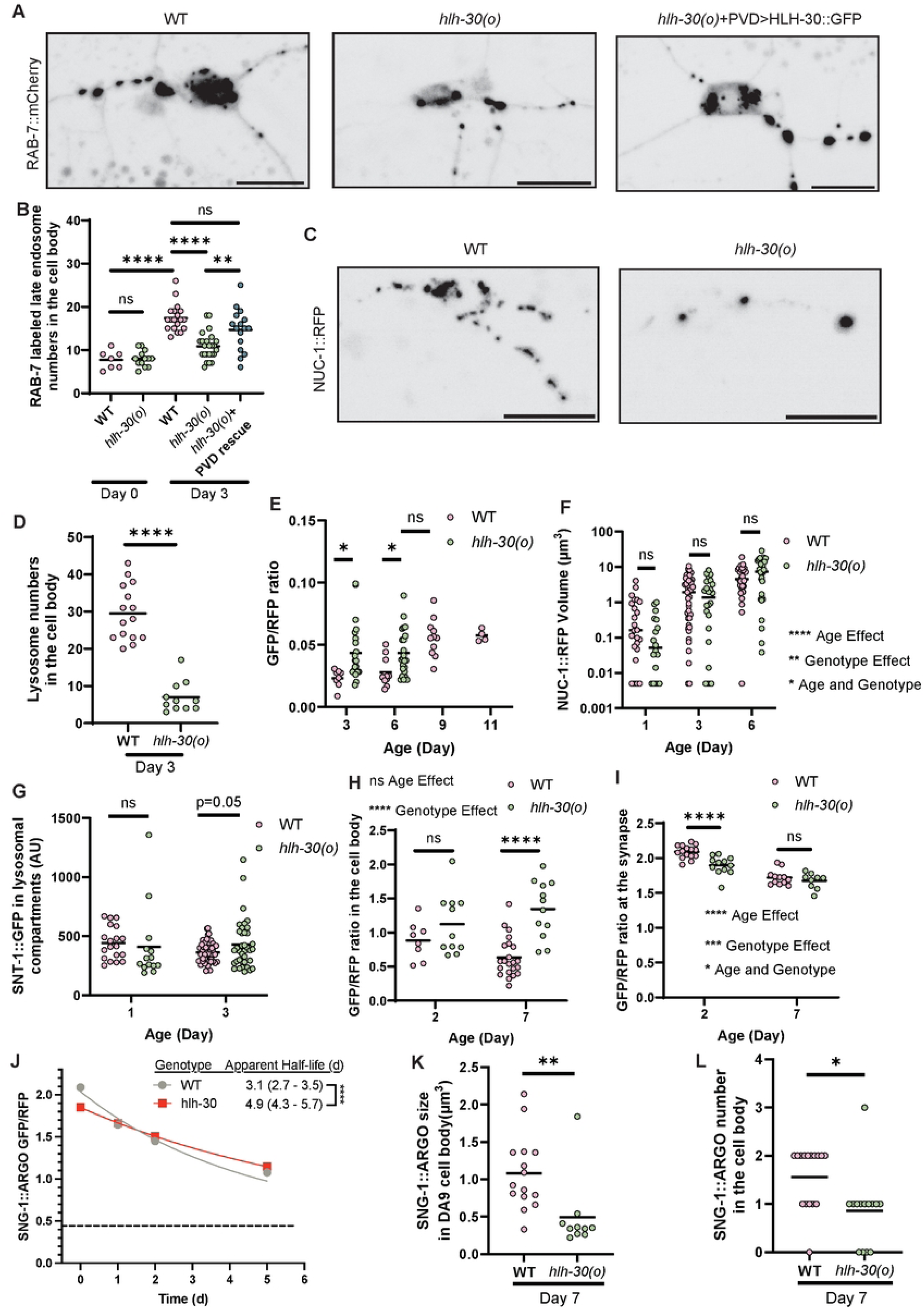
HLH-30 functions cell-intrinsically to expand neuronal lysosomal capacity in early adulthood. (A-B) The *hlh-30(o)* mutant has fewer RAB-7::mCherry-labeled endosomes in the PVD neuron cell body by Day 3 of adulthood compared to WT, and this reduction can be rescued by PVD-specific expression of HLH-30. Scale bar = 10 μm. **** P < 0.0001, **P < 0.01, ns: not significant, One-way ANOVA with Tukey post-test. (C-D) The NUC-1::RFP-positive endosome population is decreased in the *hlh-30(o)* mutant. Scale bar=10 μm. ****P < 0.0001, two-tailed *t* test. (E) The lysosomal compartment luminal pH is higher in the *hlh-30(o)* mutant by Day 3 of adulthood, similar to the aging WT phenotype. *P < 0.05, ns: not significant, One-way ANOVA with Tukey post-test. Note that the WT data is repeated from Fig 1 to facilitate comparison between the *hlh-30(o)* mutant and aging WT. (F) Neuronal lysosomal compartment size is significantly different in the *hlh-30(o)* mutant compared to WT across early adulthood (main effect of both aging and genotype). ****P < 0.0001, **P < 0.01, *P < 0.05, ns: not significant, ART ANOVA with post hoc pairwise comparisons. (G) Lysosomal degradative cargo SNT-1::GFP shows a trend of increased accumulation within lysosomal compartments in the Day 3 *hlh-30(o)* mutant compared to WT. ns: not significant, Two-way ANOVA with Sidak post-test. (H-K) Analysis of the turnover of lysosomal degradative cargo SNG-1 in the DA9 neuron using SNG-1::ARGO. Steady-state fluorescence of SNG-1::RFP::GFP was analyzed in (H, J-K); for (I), the neuron produced SNG-1::RFP::GFP until time = 0 and only SNG-1::RFP subsequently, so declining GFP shows SNG-1 protein turnover. Note that the steady-state GFP/RFP ratio at the synapses is near two rather than one because GFP is brighter than RFP. (H-I) Steady-state GFP/RFP ratio of SNG-1::ARGO in the cell body (H) and presynapses (I). The lower steady-state SNG-1::ARGO GFP/RFP ratio within vesicles in the cell body compared to at the synapses shows that the cell body vesicles are acidic lysosomal compartments. (H) The cell body SNG-1::ARGO GFP/RFP ratio increases in the *hlh-30(o)* mutant compared to WT by Day 7. (I) steady-state SNG-1::ARGO GFP/RFP ratio at the synapses is decreased in the *hlh-30(o)* mutant compared to WT at Day 2. For (H-I), **** P < 0.0001, ***P < 0.001, ns: not significant, Two-way ANOVA with Sidak post-test. (J) SNG-1::ARGO turnover rate at the synapses, calculated from one-phase exponential decay curves. **** P < 0.0001, Extra sum-of-squares F test. (K-L) The *hlh-30(o)* mutant shows decreased SNG-1::ARGO puncta size (K) and fewer puncta (L) in the neuron cell body by Day 7 compared to WT. **P < 0.01, *P < 0.05, two-tailed *t* test.

Next, we quantified NUC-1::RFP-labeled lysosomal compartments in WT versus *hlh-30(o)* mutants (Fig 2C). The number of NUC-1 labeled compartments is reduced in *hlh-30(o)* mutants, following the same trend as the RAB-7-labeled compartments population (Fig 2D). These results suggest that HLH-30 promotes the expansion of the endolysosomal system in neurons during early adulthood.

Based on these results, we hypothesized that HLH-30 is required to generate adequate lysosomal capacity in adult neurons. If this is the case, then *hlh-30(o)* mutants should exhibit accelerated neuronal lysosomal dysfunction in adulthood compared to WT. Using the PVD lysosome acidity reporter (Figs 1A-B), we found that the NUC-1::GFP/RFP ratio is higher in *hlh-30(o)* mutants compared to WT at Day 3 and Day 6 of adulthood, indicating that lysosomal compartment acidity is indeed disrupted in *hlh-30(o)* mutants (Fig 2E).

We predicted that the size of NUC-1::RFP-labeled compartments would also be larger in the *hlh-30(o)* adults, mirroring the WT aging phenotype. To test this, we compared NUC-1::RFP-labeled lysosomal compartment size between the *hlh-30(o)* mutant and WT at Day 1, 3, and 6 of adulthood. In both genotypes, the NUC-1::RFP-labeled lysosomal compartments get progressively larger with age (Fig 2F). The comparison between genotypes is more complex: at Day 1 of adulthood, there is a trend for *hlh-30(o)* mutant to have smaller mean NUC-1::RFP size compared to WT, consistent with the model that HLH-30 is necessary to promote lysosomal capacity at this age. By contrast, at Day 6 of adulthood, there is a trend for the *hlh-30(o)* mutant to have larger mean NUC-1::RFP size compared to WT, following our prediction. Day 3 of adulthood appears to be the inflection point, as NUC-1::RFP size appears no different between the genotypes. Statistically, the genotype has a significant main effect on NUC-1::RFP size when considering all ages together, and a significant difference simultaneously comparing genotype and age indicates that the lysosomal compartment size changes in *hlh-30(o)* are dependent on age, though no individual pairwise comparison between genotypes is significantly different (Fig 2F).

Next, we assessed accumulation of degradative cargo SNT-1::GFP in the lysosomal compartments, and we found an increase in the *hlh-30(o)* Day 3 adults compared to WT, driven by a subset of lysosomal compartments that exhibit more SNT-1::GFP fluorescence (Fig 2G). This is consistent with the model that lysosomal degradation is less efficient in the absence of HLH-30 function, though an alternate interpretation is that the amount of cargo that is delivered to lysosomal compartments for degradation is increased, in which case the efficiency of degradation could be unchanged.

To determine which of these models is more accurate, we set out to directly test the notion that the HLH-30-mediated expansion of lysosomal capacity is necessary for efficient turnover of degradative cargo. To do this, we applied our recently developed Analysis of Red-Green Offset (ARGO) method, wherein we endogenously tagged another characteristic cargo of lysosomal degradation, the synaptic vesicle protein Synaptogyrin/SNG-1, with both GFP and RFP (S1C Fig) [54]. The ARGO method involves a pulse-chase component in which the gene encoding *gfp* is excised via Cre/LoxP recombination with the pulse, and then the neuron is periodically imaged to quantify the ratio of GFP/RFP intensity at the synapses during the chase. A one-phase exponential decay curve was fitted to these data to calculate the presynaptic half-life of SNG-1. We induced the visualization of SNG-1 specifically in the cholinergic motor neuron Dorsal A type motor neuron 9 (DA9) using FLP/FRT recombination. We selected the DA9 neuron for this experiment because A) it allows us to investigate the turnover dynamics of a synaptic vesicle protein, and the DA9 neuron serves as a well-established model for examining presynapses; and B) we aimed to determine whether HLH-30 promotes lysosomal capacity broadly across neurons or specifically in PVD neurons.

First, we examined steady-state SNG-1::ARGO fluorescence intensity in control animals that were not “pulsed.” We observed no difference in presynapse localization, organization, or number between *hlh-30(o)* mutants and WT animals (S3B and S3C Figs). The presynaptic fluorescence intensity of SNG-1::ARGO RFP is slightly decreased in the *hlh-30(o)* mutant compared to WT, suggesting a slight reduction in synaptic vesicle numbers per synapse (S3D Fig). Consistent with the framework that endolysosomal protein degradation occurs in the neuron cell body, we observed endosomes with a lower SNG-1::ARGO GFP/RFP fluorescence intensity ratio in the cell body compared to the presynapses in WT animals (Fig 2H-I). In these cell body-localized endosomes, the GFP fluorescence from SNG-1::ARGO is partially quenched, indicative of the low pH-environment of degradative endolysosomes (Fig 2H).

Next, with the ARGO pulse at Day 2 of adulthood, we found that the SNG-1 presynaptic turnover rate is slower in *hlh-30(o)* mutants (half-life = 4.9 d, 95% C. I. 4.3-5.7 d) compared to WT (half-life = 3.1 d, 95% C. I. 2.7-3.5 d) (Fig 2J, S3E and S3F Figs). Interestingly, the SNG-1::ARGO GFP/RFP ratio is lower at the presynapses in the *hlh-30(o)* mutant compared to WT at Time = 0 of the experiment, which shows the steady-state fluorescence before the “pulse” (Fig 2I-J). Previous research indicates that synaptic vesicle proteins are sorted for degradation into late endosomes at the presynapses [7,8,55]. Our result likewise indicates that SNG-1 is sorted for degradation into acidic late endosomes at the presynapses, and they suggest that in the *hlh-30(o)* mutant, a larger proportion of presynaptic SNG-1 resides in late endosomes rather than in synaptic vesicles compared to WT. Considering this change in steady-state presynaptic SNG-1::ARGO GFP/RFP along with the slower SNG-1 turnover rate in the *hlh-30(o)* mutant, it suggests that there is a backup in the clearance of sorted-for-degradation SNG-1 from the presynapses in the *hlh-30(o)* mutant.

The slower turnover of SNG-1 in the *hlh-30(o)* mutant compared to WT motivated us to take a closer look at the steady-state fluorescence of SNG-1::ARGO-labeled compartments in the cell body, which are in the process of being degraded based on their low pH (Fig 2H-I). Notably, in the *hlh-30(o)* mutants, there are fewer and smaller SNG-1::ARGO-labeled endolysosomes in the neuron cell body, suggesting reduced lysosomal capacity (Figs 2K-L). Further, the *hlh-30(o)* mutants show a higher SNG-1::ARGO intensity ratio in the neuron cell body compared to WT, suggestive of dysfunctional endolysosomal degradation (Fig 2H).

Taken together, these results indicate that HLH-30 is required for maintaining neuronal lysosomal function in early adulthood, whereas loss of *hlh-30* leads to inadequate lysosomal degradation.

### Set-point HLH-30 activity in neurons operates without nuclear enrichment and declines in aging

During starvation and stress, TFEB/HLH-30 translocates from the cytoplasm to the nucleus [20–22,56]. The subcellular localization of TFEB has been used as a key indicator of its activity status, wherein nuclear enrichment means it is active and cytosolic enrichment means it is inactive. We therefore wondered whether neuronal HLH-30 shows enriched nuclear localization when it is functioning in well-fed, unstressed young adults to expand neuronal lysosomal capacity. We used split GFP to visualize the sub-cellular localization of endogenously tagged HLH-30::GFP11 in neurons (Fig 3A, S1D Fig) [57]. Neuronal HLH-30::GFP11 shows enriched nuclear localization upon heat stress, consistent with findings in other cell types (Figs 3A-B) [20,21]. However, HLH-30::GFP11 avoids the nucleus in the steady-state of both early-(Day 3) and mid-adult (Day 6) PVD neurons (Figs 3A-B). When the animals are starved for 1 day, neuronal HLH-30::GFP11 also avoids the nucleus (Figs 3A-B). We speculate that the PVD neurons might be protected from starvation, preventing them from experiencing or sensing it during our starvation manipulation. Recent research suggests that HLH-30 activity is not always correlated with its nuclear localization [58]. Our data suggest that, although acute stress can increase neuronal HLH-30 activation by promoting its nuclear localization, the basal, or set-point, HLH-30 activity that expands lysosomal capacity in healthy, unstressed adulthood occurs in the absence of nuclear enrichment.

**Fig 3.**
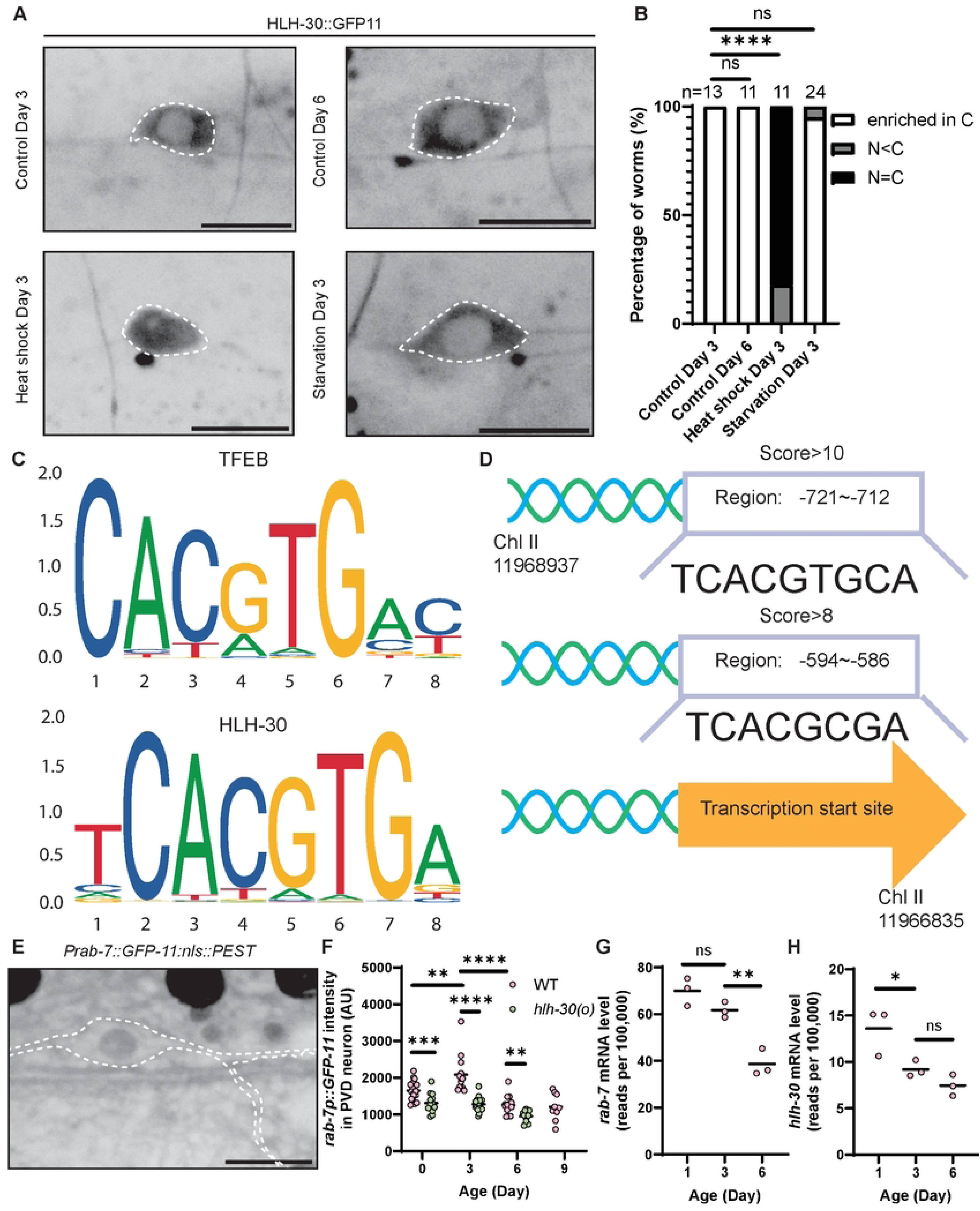
Set-point HLH-30 activity is not accompanied by enriched nuclear localization and declines during neuronal aging. (A-B) Endogenous HLH-30, visualized with split GFP, is mainly localized in the neuron’s cytoplasm in unstressed adulthood. Scale bar = 10 μm. ****P < 0.0001, ns: not significant, Fisher’s exact test. (C) The bHLH binding sequence is similar between human TFEB and *C. elegans* HLH-30 [59]. (D) High score of HLH-30 binding sequences is found in the *rab-7* promoter-plus-enhancer region. (E-F) A fluorescent reporter of *rab-7* transcription shows that PVD-specific *rab-7* transcription increases in early-adults (Day 3) and decreases during aging in WT. In *hlh-30(o)*, the PVD-specific *rab-7* transcription level is lower compared to WT at the same age, and there is no difference between Days 0, 3, and 6. Scale bar = 10 μm. The outline of the PVD neuron was generated using the PVD>bfp fluorescence in this strain, which is not shown. **** P < 0.0001, ***P < 0.001, **P < 0.01, One-way ANOVA with Tukey post-test. (G) Relative *rab-7* mRNA abundance from whole-worm transcriptomic data shows reduced *rab-7* mRNA level in aging WT worms. **P < 0.01, ns: not significant, One-way ANOVA with Tukey post-test [60]. (H) Relative *hlh-30* mRNA abundance from whole-worm transcriptomic data shows reduced *hlh-30* mRNA level in aging WT worms. *P < 0.05, ns: not significant, One-way ANOVA with Tukey post-test [60].

We hypothesized that basal HLH-30 activity declines in aged neurons, contributing to the aging-associated reduced lysosomal function. To test this hypothesis, we generated a real-time fluorescent transcriptional reporter of *rab-7*, wherein we express *GFP11::nls::PEST* from the endogenous promoter-plus-enhancers of *rab-7* and use neuronally-expressed *GFP1-10* to reconstitute GFP fluorescence specifically in the neurons (S1E Fig). *RAB7* is a known transcriptional target of TFEB in mammalian cells [12]. HLH-30 shares a similar bHLH binding sequence as TFEB, and there are several predicted binding sites of HLH-30 in the promoter and enhancer region of *rab-7* (Figs 3C-D) [59]. The *GFP11::nls::PEST* is transcribed together with *rab-7* into the same mRNA, which is then alternatively spliced to separate the *GFP11::nls::PEST* mRNA from the *rab-7* mRNA, to be translated separately. The PEST sequence targets protein for rapid degradation, so the fluorescence intensity of GFP11 reports the transcription level of *rab-7* within a short period, instead of the cumulative transcription level [60]. The NLS (nuclear localization signal) directs GFP11 to the nucleus, facilitating image quantification [61]. The transcription level of *rab-7* in WT PVD neurons increases in early adulthood (Day 3), corresponding with the expansion of the number of late endosome and lysosomal compartments. Then, it decreases in older neurons (Day 6), anticipating the age-associated decline of lysosomal function (Fig 3E-F). In *hlh-30(o)* mutants, the transcription level of *rab-7* in the neuron is always lower compared to WT at the same age (Fig 3F). Furthermore, in contrast to WT animals, we detected no difference in *rab-7* transcription level between Day 0, Day 3, and Day 6 adults in *hlh-30(o)* mutants (Fig 3F).

We analyzed pre-existing RNAseq data from whole animals to assess whether the changes we observed in neuronal *rab-7* transcription during adulthood extend to the whole animal [62]. The whole-animal *rab-7* mRNA level decreases from Day 3 to Day 6, consistent with the decrease observed with our neuron-specific transcriptional reporter (Fig 3G). On the other hand, there is no increase in the *rab-7* mRNA level in early adulthood, suggesting that the expansion of lysosomal capacity during early adulthood may be neuron-specific. We next examined the mRNA levels of five additional genes, *cpr-1*, *cpr-5*, *vha-5*, *lmp-2*, and *syx-17*, that are well-validated transcriptional targets of HLH-30 [63]. mRNA levels of *cpr-5*, *lmp-2*, and *syx-17* show a statistically significant decrease during aging. Therefore, 4 out of 6 HLH-30 transcriptional targets show reduced expression in aging, by bootstrapping analysis, only 3.5% of randomly selected groups of 6 genes have at least 4 that decline in aging (S4 A-G Figs). Moreover, the level of *hlh-30* mRNA itself also decreases during aging (Fig 3H). HLH-30 is predicted to regulate its own expression based on ChIP-seq data [64], echoing the self-transcriptional regulation of mammalian TFEB [65]. Together, these results suggest that HLH-30 transcriptional activity decreases with age, likely contributing to the age-associated neuronal lysosomal dysfunction.

### Loss of HLH-30 accelerates age-related decline in dendrite morphology maintenance

In WT worms, the PVD neuron displays a highly branched and organized dendritic arbor. The structure is developed by Day 0, and the PVD neuron can fully maintain organization in middle-aged adult, Day 6 worms (Fig 4A). However, in Day 6 *hlh-30(o)* mutants, we observed disorganized dendrite outgrowths, what we refer to as the “sprouting” phenotype, and dendrite degeneration, in approximately 75% and 50% of the population, respectively (Figs 4A-C). We did not notice any dendrite degeneration or sprouting in either WT or *hlh-30(o)* at Day 0 and Day 3 (S2 Fig). Previous works have shown that the PVD dendrite exhibits progressive degeneration in WT as they age [66,67]. We likewise observed progressive dendrite degeneration with age in WT animals, and the *hlh-30(o)* mutant at Day 6 exhibits a similar level of dendrite degeneration as WT at Day 11 (Fig 4B). We also observed that the sprouting phenotype increases with age in WT animals, and the percentage of animals showing sprouting at Day 11 in WT is similar to that in *hlh-30(o)* at Day 6 (Fig 4C). These results suggest that loss of HLH-30 causes early-onset and accelerated neuronal aging in PVD neurons. Expression of wild-type *hlh-30* from a pan-somatic promoter was able to rescue the level of both sprouting and dendrite degeneration in *hlh-30(o)* to those observed in WT at Day 6, confirming that HLH-30 is necessary for maintaining dendrite morphology in aging (Figs 4 D-E).

**Fig 4.**
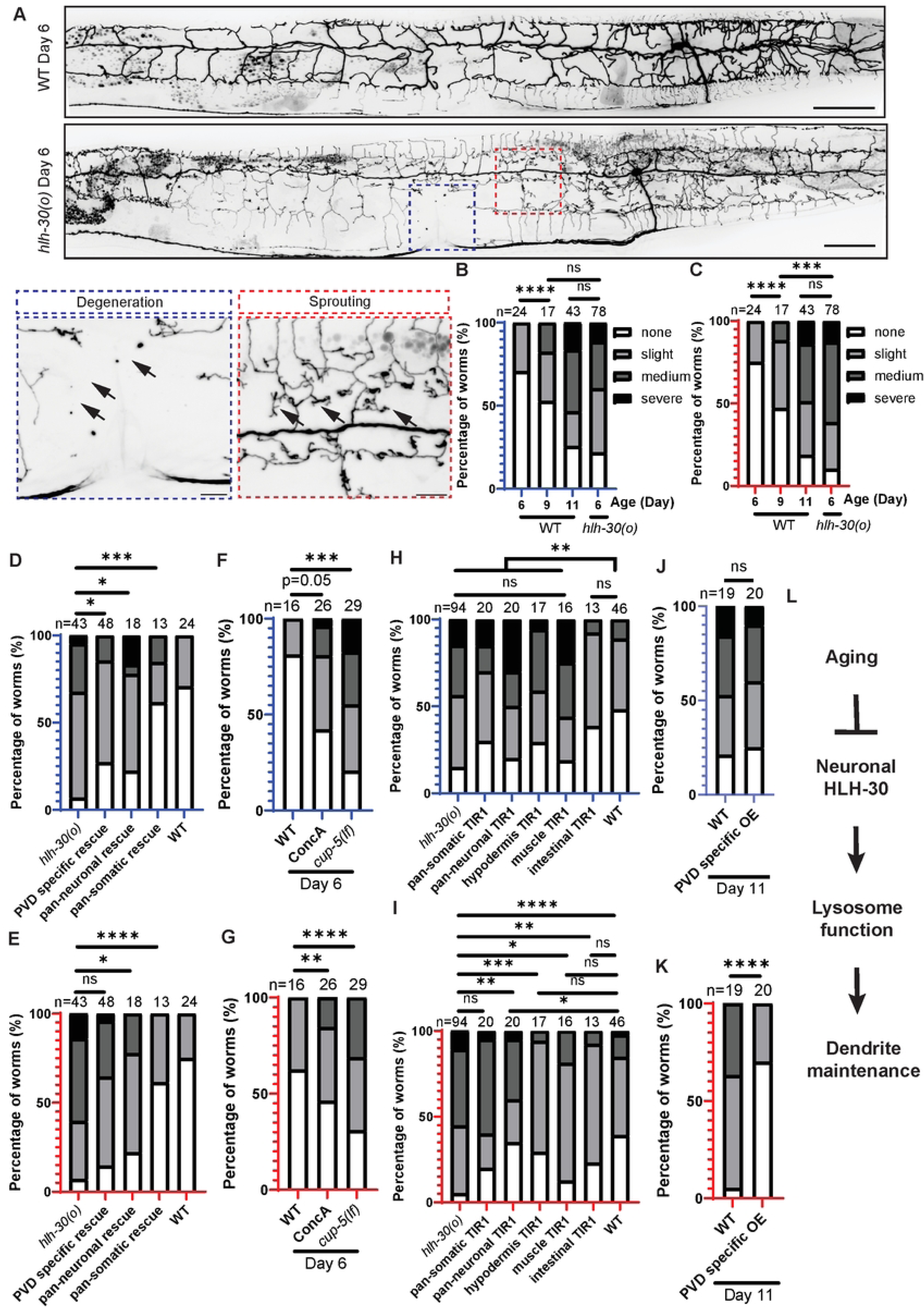
Loss of HLH-30 accelerates age-related dendrite degeneration and sprouting. (A-C) In WT adults, the PVD dendrite shows progressive dendrite degeneration (B) and sprouting (C) with age, and these phenotypes manifest earlier in the *hlh-30(o)* mutant. Note that WT Day 11 shows similar penetrance and severity as *hlh-30(o)* mutant Day 6. In (A), the blue and red dashed boxes on the *hlh-30(o)* Day 6 animal, which correspond to the zoomed-in images below with the same outline colors, show examples of severe dendrite degeneration and severe sprouting, respectively. Scale bar = 50 μm (whole worm) or 10 μm (zoom in). ****P < 0.0001, ***P < 0.001, ns: not significant, Fisher’s exact test. Arrowheads point to degeneration or sprouting (not all instances are indicated). (D-E) Pan-somatic expression of *hlh-30* rescues the dendrite degeneration (D) and sprouting (E) phenotypes in Day 6 *hlh-30(o)* mutants, and PVD-specific or pan-neuronal *hlh-30* expression partially rescues the dendrite defects. ****P < 0.0001, ***P < 0.001, *P < 0.05, ns: not significant, Fisher’s exact test. (F-G) WT animals exposed to Concanamycin A treatment and *cup-5(lf)* mutant animals both show increased percentage and severity of dendrite degeneration (F) and sprouting (G) compared to WT at Day 6. ****P < 0.0001, ***P < 0.001, **P < 0.01, ns: not significant, Fisher’s exact test. (H-I) Depleting HLH-30 in the whole worm (except the germline) phenocopies the *hlh-30(o)* mutant, showing increased dendrite degeneration (H) and sprouting (I) compared to WT at the same age. Note that for these experiments, all animals were cultivated in the presence of Auxin, and they were scored at a later age than A-G because the Auxin caused slowed growth and aging. (H) Pan-neuronal, muscle, and hypodermal-specific HLH-30 depletion all show increased degeneration compared to WT. **P < 0.01, ns: not significant, Fisher’s exact test. (I) Pan-neuronal HLH-30 depletion causes increased sprouting compared to WT. ***P < 0.001, **P < 0.01, *P < 0.05, ns: not significant, Fisher’s exact test. (J-K) PVD-specific overexpression of *hlh-30* in the WT genetic background reduces severity of aging-associated sprouting (K) and does not have a significant impact on dendrite degeneration (J). ****P < 0.0001, ns: not significant, Fisher’s exact test. (L) Model for HLH-30 regulation of dendrite maintenance during aging.

Both TFEB and HLH-30 regulate genes that are not involved in the autophagy or lysosomal pathway [63,68–70]. Given our results showing that HLH-30 promotes lysosomal capacity in *C. elegans* neurons (Figs 2-3), we hypothesized that both the dendrite sprouting and degeneration phenotypes result from decline of lysosomal function. To test this, we examined the PVD dendrite morphology in WT worms treated with V-ATPase inhibitor Concanamycin A, which causes reduced acidity of lysosomal compartments (Fig 1B). Indeed, WT worms treated with Concanamycin A show an increased percentage and severity of both sprouting and degeneration compared with controls (Figs 4F-G). We also assessed the PVD dendrite morphology in *cup-5(lf)* mutant animals, which are known to have disrupted lysosomal functions [71,72]. Indeed, *cup-5(lf)* phenocopies *hlh-30(o)* by showing the sprouting and dendrite degeneration phenotypes at Day 6 (Figs 4F-G). These results indicate that both dendrite sprouting and degeneration are caused by decline of lysosomal function.

To determine the spatial requirement for HLH-30 to preserve dendrite morphology during aging, we used the Auxin-inducible degradation (AID) system, wherein we inserted the *AID* sequence into the *hlh-30* genomic locus to tag the C-terminus of HLH-30 (S1F Fig). When the hormone Auxin is added, the tissue-specifically expressed TIR1 E3 ubiquitin ligase recognizes the AID-tagged HLH-30 and promotes its degradation [73]. This technique allows us to deplete HLH-30 in specific tissues. Auxin delays worm growth, and the progression of age-related dendrite phenotypes irrespective of the presence of TIR1, so we performed all comparisons for this experiment in Day 9 adults grown on Auxin (Figs 4H-I). We used pan-somatically expressed TIR1 to eliminate HLH-30 in the whole worm except the germline, which phenocopies the dendrite degeneration and sprouting phenotypes observed in the *hlh-30(o)* mutant (Figs 4H-I). Next, we depleted HLH-30 from four individual somatic tissues: neurons, hypodermis, muscle, and intestine. Neuron, hypodermis and muscle-specific degradations all show a significant increase in dendrite degeneration compared to WT, and the severity is similar to that of the *hlh-30(o)* mutant (Fig 4H). This suggests that HLH-30 is required in neurons, as well as in the two tissues that physically contact the PVD dendrite, to protect against dendrite degeneration. For the dendrite sprouting phenotype, pan-neuronal HLH-30 depletion shows an intermediate sprouting phenotype between *hlh-30(o)* and WT (Fig 4I). Muscle-specific HLH-30 degradation also results in a trend of higher percentage of the population exhibiting sprouting compared to WT, though it is not statistically significant (Fig 4I). These data indicate that HLH-30 is necessary in neurons and likely also in muscle to maintain organized PVD dendrites during aging.

Since HLH-30 functions cell-autonomously to expand lysosomal compartment abundance (Figs 2A-B), we asked whether HLH-30 in the PVD neuron is likewise sufficient to protect against age-associated dendrite morphology defects. Indeed, overexpressing HLH-30 either specifically in PVD neurons or pan-neuronally can partially rescue the sprouting and dendrite degeneration phenotypes in the Day 6 *hlh-30(o)* mutant (Figs 4D-E). These results are consistent with the results from the AID experiment (Figs 4H-I), and they indicate that HLH-30 is working in both cell-autonomous and non-cell-autonomous pathways to preserve adult dendrite morphology. The *hlh-30(o)* mutant is reported to have a wild-type lifespan at 20 °C and a slightly shortened lifespan at 25 °C [19,74]. Importantly, the shortened lifespan at 25 °C cannot be rescued with neuron-specific HLH-30 expression [74]. Therefore, the rescue of the *hlh-30(o)* mutant’s dendrite degeneration and sprouting by PVD-specific or pan-neuronal *hlh-30* expression is not an indirect effect of lifespan differences between the strains.

Furthermore, PVD-specific overexpression of *hlh-30* in WT worms reduces the severity of sprouting at Day 11, though it has no statistically significant impact on dendrite degeneration (Figs 4J-K). These data indicate that HLH-30 is sufficient to partially protect against age-associated dendrite morphology defects, supporting a model in which the basal, or set-point activity of HLH-30 during adult homeostasis is essential for preserving lysosomal function and thereby maintaining neuron integrity during aging.

## Discussion

Here, we established a vital role for HLH-30/TFEB in maintaining lysosomal homeostasis in mature neurons *in vivo*. We proposed a model in which HLH-30/TFEB operates intrinsically within neurons of healthy young adults, in the absence of starvation or exogenous stress, to expand lysosomal capacity (Fig 4L). This likely aids in remediating the rising proteostasis demand during aging. In contrast to the stress conditions that promote HLH-30 translocation to the nucleus, HLH-30 functions in basal homeostasis while predominantly enriched in the cytoplasm. Our model further proposes that HLH-30 activity declines in aged adults, causing reduction of neuronal lysosomal degradative capacity, which in turn contributes to declining maintenance of dendrite morphology (Fig 4L).

We demonstrated that neuronal lysosomal function declines with age in healthy, wild-type animals based on three established features of lysosome dysfunction: 1) failure to maintain the acidic lysosomal pH, 2) enlarged lysosomal compartments, and 3) lysosomal accumulation of degradative cargo (Fig 1) [44]. This decline in lysosomal function is consistent with previous findings across multiple species, including humans, mice, *D. melanogaster,* and *C. elegans,* showing that autophagic activity becomes impaired in aging neurons, potentiating the accumulation of neurodegenerative disease-associated protein aggregates and defective organelles [36,75–78]. Given that autophagic flux requires efficient lysosomal degradation, we propose that neuronal lysosomal dysfunction partially underlies this age-associated decline in autophagy. Importantly, comparing our finding to previous works, lysosomal dysfunction appears to precede other cellular pathologies of neuron aging, including synapse loss and axon morphology defects (Fig 1) [79–81]. This sequence of events suggests that lysosomal dysfunction is an early molecular event in neuronal aging, likely contributing to impaired autophagic flux and declining proteostasis [82].

TFEB is well-established as a master regulator of lysosome biogenesis, especially in response to starvation [11,13,14]. Its potential to enhance the autophagic clearance of pathogenic accumulations of lysosomal degradative cargo marks TFEB as a compelling therapeutic target for neurodegenerative disease. Indeed, over-expression of TFEB has demonstrated benefits in multiple experimental models of Alzheimer and Parkinson’s disease [83–88]. Our findings expand the framework for thinking about TFEB and neurodegenerative disease, suggesting that it is not only a potential therapeutic target but also a plausible underlying mechanism. Specifically, evolutionary conservation of the *C. elegans* function of HLH-30 would implicate TFEB in promoting adult neuron maintenance, and its declining activity in contributing to age-associated lysosomal dysfunction that may potentiate neurodegenerative disease [89]. Supporting the notion that this role in neuron homeostasis may be conserved, knockdown of TFEB/Mitf in *Drosophila* neurons results in an apparent blockage of autophagic flux in the adult fly brain in the absence of starvation or stress [90]. Further, brain-specific TFEB knockout in mice leads to increased Aβ and Tau protein accumulation, apoptotic cells, and axonal degeneration; it would be interesting to assess to what extent these phenotypes are due to loss of TFEB in neurons versus in glial cells [91,92].

It is generally accepted that when TFEB/HLH-30 is active, it shows a corresponding enriched nuclear localization [13,14,89,93]. However, our data indicate that HLH-30 promotes neuronal lysosomal capacity during steady-state maintenance while exhibiting enriched cytoplasmic localization (Figs 3A-B, Figs 3F-G, S3 Figs). Additionally, we observed reduced HLH-30 transcriptional output during aging without a corresponding change in its subcellular localization, further decoupling HLH-30 function from its cytosolic versus nuclear subcellular localization in the context of basal homeostasis. This result corroborates recent research suggesting that HLH-30 nuclear localization is not necessarily strongly correlated with its transcriptional output [58]. It would be interesting to investigate whether the transcriptional output of neuronal HLH-30 changes in composition as well as in strength during basal homeostasis versus stress [21]. Moreover, our results raise the consideration that a lack of nuclear enrichment for mammalian TFEB may not be a strong indicator that it does not have an important function in promoting a low level of basal transcription in a given context.

Our data indicate that neuron-intrinsic lysosomal dysfunction leads to defective maintenance of dendrite morphology, causing both dendrite degeneration and sprouting. What mechanism links these phenotypes? For dendrite degeneration, one possibility is that lysosomal dysfunction causes a buildup of autophagosomes in the dendrite. These could locally disrupt transport or function of necessary cellular components, leading to degeneration. Indeed, autophagosomes are observed within the beading or swelling structures that are early signs of degeneration, and formation of autophagosomes within the neuron promotes dendrite degeneration [66]. A decline in autophagic flux might also lead to the accumulation of damaged mitochondria, increasing oxidative stress or disrupting calcium homeostasis that could promote neurodegeneration [94]. Alternatively, or in addition, an accumulation of unwanted protein cargoes of endolysosomal degradation in the dendrite could disrupt structural integrity or ion homeostasis [95]. For dendrite sprouting, endocytic trafficking of dendrite guidance receptors is critical for maintaining proper receptor activity at the plasma membrane, so it is possible that disrupted lysosomal function leads to guidance receptor buildup on the plasma membrane, leading to the sprouting phenotype[96–98]. Interestingly, our results indicate that HLH-30 promotes dendrite maintenance both cell-intrinsically and non-cell-autonomously (Fig 4), so some extracellular pathways are also involved in both dendrite degeneration and sprouting. The observation that the *cup-5(lf)* mutant phenocopies *hlh-30(o)* in regard to dendrite degeneration but exhibits less severe sprouting suggests that additional mechanisms beyond lysosomal defects may contribute to the sprouting phenotype (Figs 4F-G). TFEB and HLH-30 target genes involved in lysosomal biogenesis and autophagy are only a subset of their transcriptional targets; others include genes involved in metabolism, mitochondrial homeostasis, and protein transport [99,100]. Indeed, lysosome-independent functions of TFEB include regulating lipid metabolism during starvation in the liver, glucose homeostasis during exercise in the muscle, inflammatory response in macrophages, and apoptosis in cancer cells [65,68,101,102]. Loss of one of the lysosome-independent function of HLH-30 may contribute to the dendrite sprouting phenotype.

Inadequate lysosomal degradation is expected to cause a buildup in degradative cargo within lysosomes [44]. Indeed, both aging and loss of HLH-30 lead to accumulation of lysosomal cargo Synaptotagmin/SNT-1 within lysosomal compartments (Figs 1 and 2). Further, we found that this buildup extends to the presynapse. Specifically, our data suggest that in the *hlh-30(o)* mutant, the presynaptic half-life of SNG-1 is prolonged compared to WT, and this is accompanied by an increase in the proportion of presynaptic SNG-1 that resides in acidic late endosomes versus synaptic vesicles. This indicates that while SNG-1 is sorted into acidic late endosomes at the synapse in both WT and the *hlh-30(o)* mutant, the clearance of these degradation-targeted SNG-1 molecules is delayed in the *hlh-30(o)* mutant (Figs 2H-I). Our findings are consistent with prior studies in *Drosophila* and mammalian neurons indicating that synaptic vesicle proteins are sorted for degradation at the presynapse, and they suggest that lysosome functionality contributes to regulating the half-life of presynaptic proteins [7,8,55,103]. Together, these findings suggest a mechanism wherein lysosome dysfunction causes impaired removal of presynaptic degradative cargoes, which could contribute to neuronal dysfunction.

In summary, understanding how neurons adapt to proteostasis demands throughout aging is crucial for uncovering the mechanisms that support neuronal maintenance. Our work identifies HLH-30 as a key factor that expands lysosomal capacity during young adulthood. Moreover, our findings suggest that the set-point level of HLH-30 activity determines, in part, the rate of age-associated lysosomal dysfunction and subsequent decline in dendrite maintenance. Considering this together with the demonstrated therapeutic potential of TFEB activation in neurodegenerative disease models, provides a strong foundation for future therapeutic strategies to stave off neuron aging and dysfunction [84,85,88].

## Materials and methods

### *C. elegans* strains and maintenance

For maintenance, *C. elegans* strains were grown on nematode growth medium (NGM) seeded with *E. coli* OP50 at room temperature (22 ℃) [104]. Aged worms are defined by the number of days after the L4 stage, “Day 0” of adulthood. All experiments were performed at room temperature (22 ℃), except for those in Figs 1C-E, Figs 2F-L, and S3B-F Figs, which were performed at 20 °C. Wild-type N2 and JIN1375 *hlh-30(tm1978) IV* strains were obtained from the Caenorhabditis Genetics Center (CGC), which is funded by NIH Office of Research Infrastructure Programs (P40 OD010440). Strains used in this study are listed in S1 Table.

### Cloning and strain generation

Cloning was carried out in the pSM vector, a derivative of pPD49.26, or pPD117.01 (Addgene, Watertown, MA, USA) using standard restriction enzyme cloning technique (NEB, *Ipswich, MA, USA*). *Podr-1::gfp*, *Podr-1::rfp*, *Pdes-2::bfp*, and *Punc-122::rfp* vectors are kindly provided by the Kang Shen lab, Stanford University. *nuc-1, hlh-30,* and *rab-7* cDNA were amplified from a cDNA prepared from total RNA isolated from N2 worms. Sequence of the primers are listed in S2 Table. Transgenes expressed from extrachromosomal arrays or integrated arrays were generated by microinjection of the constructs [105]. Genome editing was carried out by CRISPR-Cas9 [106]. The *hlh-30 syb* allele was generated at SunyBiotech (Fuzhou, Fujian, China) by CRISPR-Cas9.

### Confocal imaging and fluorescence microscopy

Worms were transferred to a slide with a 3% agarose pad made by melting agarose in M9 buffer. Worms were immobilized in 10 mM sodium azide in M9 buffer for experiments to examine HLH-30::GFP localization and 20 mM levamisole in M9 buffer for all other experiments. Images were captured using a Nikon eclipse Ti2 microscope paired with a CSU W1 SoRa confocal scanner unit and a YOKOGAWA spinning disc unit with a Plan Apo VC 60xA/1.20 WI objective. Image settings (laser power and exposure time) were identical for all genotypes and treatment groups across experiments. Images were taken with 0.5 μm step size in z-stack for 10-25 μm range. Images were analyzed using ImageJ software. Fluorescence intensities and sizes of vesicles were quantified from images using “3D object counter” (Figs 1B and 1E, Figs 2E-K, Fig 3F, S3D Fig). Mean fluorescence intensities were used for all figures and calculations except for *rab-7* transcription level (Fig 3F), which used maximum fluorescence intensities. Number of vesicles were quantified from images using “3D object counter” (Fig 1C, Fig 2L), by hand (Figs 2B and 2D, S3A Fig) or using “find maxima” (S3C Fig). HLH-30 localizations were quantified from images by eye. PVD dendrite morphology were quantified by eye using the Nikon eclipse compound microscope.

For experiments using SNG-1::ARGO, maximum projection images were generated from z-stack of the whole presynaptic region of each animal. Presynapses were identified by generating a mask from the RFP image with default threshold settings followed by watershed. This mask was used to calculate the mean fluorescence intensity of each synapse in both the RFP and GFP images. The GFP/RFP was calculated for each presynapse detected, then averaged to generate one presynaptic GFP/RFP value for each animal.

Images of HLH-30 localization were taken in the green channel (Image setting: 100 ms exposure time, 60% laser power). The look up table (LUT) of green channel was adjusted to 30-800 in ImageJ before quantifying. Categories of HLH-30 localization were performed as follow: neurons with HLH-30::GFP enriched in the cytoplasm and has a clear boundary between nucleus and cytoplasm were counted as enriched in the cytoplasm (enrich in C); neurons with HLH-30::GFP intensity in the cytoplasm that is higher than in the nucleus, but no clear boundary between nucleus and cytoplasm were counted as cytoplasm higher than nucleus (N<C); neurons with no difference in HLH-30::GFP intensity between nucleus and cytoplasm were counted as nucleus equal to cytoplasm (N=C).

Quantification of dendrite degeneration and sprouting was performed blind to genotype/condition.

To quantify dendrite degeneration, one dendrite from each worm was assigned to a category as follows: worms with dendrite that shows several swellings and the dendrite between swellings are relatively thinner than normal dendrites were counted as “slight dendrite degeneration;” worms with dendrite that have swellings that are disconnected from the main part of the dendrite were counted as “medium dendrite degeneration;” worms with dendrite that have more than 5 branches that are disconnected from the main part of the dendrite were counted as “severe dendrite generation;” worms without any feature mentioned above were counted as “no dendrite generation.” Worms were always categorized by the most severe feature found in the whole dendrite.

To quantify dendrite sprouting, one dendrite from each worm was assigned to a category as follows: worms with disorganized dendrite outgrowth only at quaternary dendrites were counted as “slightly sprouting;” worms with disorganized dendrite outgrowth not only at quaternary dendrites, but restricted to one or two small areas were counted as “medium sprouting;” worms with disorganized dendrite outgrowth not only at quaternary dendrites among the whole dendrite were counted as “severe sprouting;” worms without any feature mentioned above were counted as “no sprouting.”

### Concanamycin A treatment

NGM plates were supplemented to a final concentration of 0.03 μM Concanamycin A (Chemcruz) from a 0.5 mM stock solution. Worms were moved onto seeded Concanamycin A plates at the L4 stage/Day 0 of adulthood and were counted or imaged at Day 6 of adulthood.

### Auxin treatment

NGM plates were supplemented to a final concentration of 1 mM K-NAA (PhytoTech Labs) from a 250 mM stock solution. Worms were synchronized by egg laying 15 gravid hermaphrodites for 2 hours on seeded Auxin plates, and PVD dendrite morphology were counted at Day 9 unless otherwise noted.

### Heat shock treatment

For measuring HLH-30 localization under heat stress, worms growing at room temperature were shifted to 35 ℃ for 3 hours at Day 3 of adulthood and were imaged immediately afterward [21]. For protein turnover experiments using SNG-1::ARGO, worms growing at 20 ℃ were shifted to 34 ℃ for 1 hour at Day 2 of adulthood, returned to 20 ℃, and then imaged 0 - 5 days afterward [54].

### Starvation treatment

Well-fed worms were moved to unseeded NGM plates at Day 2 of adulthood and were imaged at Day 3 adulthood.

### RNAseq data analysis

RNAseq data were obtained from NCBI’s Gene Expression Omnibus. Sample “SRR19895471”, “SRR19895472”, “SRR19895473”, “SRR19895474”, “SRR19895475”, “SRR19895476”, “SRR19895477”, “SRR19895478”, and “SRR19895479” from project “PRJNA853940” were downloaded from NCBI [60]. Fastq reads were first trimmed and filtered for quality using TrimGalore (0.6.10). Reference genome were build using hisat2-build based on *C. elegans* Bristol N2 DNA reference. Each sample was aligned to the reference using hisat2 (2.1.0) and the results were organized using samtools sort (1.10). The raw gene hits were quantified using featureCounts (1.6.4). Raw gene hit count tables were normalized to total number of transcripts.

### Statistical analysis

Statistical analyses were performed in either Microsoft Excel or GraphPad Prism 9. For quantitative data, single parametric pairwise comparisons were analyzed using unpaired two-tailed Student’s *t* test, and non-parametric pairwise comparisons were analyzed using Kolmogorov-Smirnov test. Multiple parametric comparisons were analyzed using one-way analysis of variance (ANOVA) test with Tukey post-test or two-way ANOVA with Sidak post-test as indicated in the figure captions. For Fig 2F, we conducted an Aligned Rank Transform (ART) ANOVA to assess the effects of genotype and age on NUC-1::RFP-labeled lysosomal compartment size. This non-parametric approach was used to accommodate the non-normal distribution of our data. The ART ANOVA was performed using the ARTool package in R, with subsequent post hoc pairwise comparisons conducted using the art.con() function. For qualitative data, Fisher’s exact test was performed between each individual comparison, and Bonferroni correction were used to determine the significance threshold for the P value. To compare SNG-1::ARGO half-lives between WT and *hlh-30(o)*, Prism 10 was used to generate one phase exponential decay curves, constraining the plateau to the experimentally determined value based on background GFP fluorescence (which comes from autofluorescence from the worm cuticle). The curves were fitted using least squares regression with no weighting, and the comparison between WT versus *hlh-30(o)* was performed using Extra sum-of-squares F test.

### Bootstrapping

200 groups of 6 genes per group were randomly chosen from the RNAseq raw gene hits list. Raw gene hit count tables were normalized to total number of transcripts.

## Acknowledgments

Some *C. elegans* strains were provided by the Caenorhabditis Genetics Center (CGC) and the Kang Shen lab, Stanford University. We thank Sophie Baumberger for generating strains, Adam Darlington for data analysis, Sherlyn Wijaya, Manuel Alvarez, Namita Khajanchi, Jiarong Gao and Jingting Liang for feedbacks on the manuscript, and members of the Richardson lab for discussions.

## Supporting information

**S1 Fig. Schematics showing the design of transgenes and alleles used.** (A) Lysosome acidity reporter, composed of three co-injected plasmids. (B) Lysosome degradative capacity reporter. The *nuc-1::rfp* and *bfp* morphology marker were co-injected, generating the transgene *carEx4.* The *Pnhr-81>Flippase* expresses in seam cells, including the PVD neuron’s grandmother, and was expressed from *wyIs836* [107]. The *snt-1(ox698)* allele is FLP-on SNT-1::GFP [108]. (C) Design for the SNG-1::ARGO-tag [54]. (D) Design for fluorescent reporter of endogenous HLH-30 localization. (E) Design for *rab-7* transcriptional reporter, which was inserted into the *rab-7* endogenous genomic locus directly after the start codon. (F) Design of the *hlh-30::AID* allele and tissue-specific TIR1 expression transgenes for the Auxin inducible degradation (AID) experiments.

**S2 Fig. PVD neuron dendrite morphology is indistinguishable between WT and the *hlh-30(o*) mutant at Day 0 and Day 3 of adulthood.** Scale bar = 50 μm.

**S3 Fig.** (A) The *hlh-30(o)* mutant has fewer endogenously tagged GFP::RAB-7::GFP endosomes in the PVD neuron cell body at Day 3 of adulthood compared to WT, and shows no difference at Day 0. **P < 0.01, ns: not significant, Two-way ANOVA with Sidak post-test. (B) Line scan images of the RFP channel in DA9 neuron synapses labeled by SNG-1::ARGO shows no apparent difference in synapse organization in the *hlh-30(o)* mutant compared to WT. Scale bar = 10 μm. (C-D) The number of DA9 presynapses shows no apparent difference between WT and the *hlh-30(o)* mutant (C), but the SNG-1::ARGO RFP fluorescence intensity is slightly decreased in *hlh-30(o)* compared to WT (D). ***P < 0.001, **P < 0.01, ns: not significant, Two-way ANOVA with Sidak post-test. (E-F) Quantification of SNG-1::ARGO GFP/RFP ratio at the synapse after the heat shock “pulse” in WT (E) and the *hlh-30(o)* mutant (F). Each data point shows the average GFP/RFP ratio at each presynapse for one animal. These data were used to generate the decay curves for Fig 2J.

**S4 Fig. mRNA abundance of HLH-30 target genes decreases with aging.** (A-E) In addition to *rab-7* (Fig 3G), three out of five more HLH-30-regulated genes show reduced mRNA level along with aging. **P < 0.01, *P < 0.05, ns: not significant, One-way ANOVA with Tukey post-test. [60] (F) Bootstrapping analysis showing numbers of genes in randomly selected groups of 6 genes that have reduced mRNA level with aging.

**S1 Table. *C. elegans* strains used in this paper.**

**S2 Table. Primers used in this paper.**

